# Using TCR and BCR sequencing to unravel the role of T and B cells in abdominal aortic aneurysm

**DOI:** 10.1101/2022.12.01.518788

**Authors:** Christin Elster, Miriam Ommer-Bläsius, Alexander Lang, Tanja Vajen, Susanne Pfeiler, Milena Feige, Khang Lê Quý, Maria Chernigovskaya, Malte Kelm, Holger Winkels, Susanne Schmidt, Victor Greiff, Norbert Gerdes

**Author notes:** Corresponding authors: Norbert Gerdes, PhD, Division of Cardiology, Pulmonology and Vascular Medicine, Medical Faculty and University Hospital, Heinrich-Heine University, Moorenstr. 5, 40225 Düsseldorf, Victor Greiff, Department of Immunology, Institute of Clinical Medicine, Rikshospitalet, Sognsvannsveien 20, 0372 Oslo.

## Abstract

**Background:** Abdominal aortic aneurysm (AAA) is a life-threatening cardiovascular disease, and the pathogenesis is still poorly understood. Recent evidence suggests that AAA displays characteristics of an autoimmune disease and it gained increasing prominence that specific antigen-driven T cells in the aortic tissue may contribute to the initial immune response. Single-cell RNA T- and B cell receptor (TCR and BCR) sequencing is a powerful tool to investigate TCR and BCR clonality and thus to further test this hypothesis. However, difficulties such as very limited numbers of isolated cells must be considered during implementation and data analysis making biological interpretation of the data challenging. Here, we perform a representative analysis of scRNA TCR and BCR sequencing data of experimental murine AAA and show a reliable and streamlined bioinformatic processing pipeline highlighting opportunities and limitations of this approach.

**Methods:** We performed single-cell RNA TCR and BCR sequencing of isolated lymphocytes from the infrarenal aortic segment of male C57BL/6J mice 3, 7, 14, and 28 days after AAA induction via elastase perfusion of the aorta. Sham operated mice at day 3 and 28 as well as non-operated mice served as controls.

**Results:** Comparison of complementarity-determining region (CDR3) length distribution of 179 B cells and 796 T cells revealed no differences between AAA and control nor between the disease stages. We found no clonal expansion of B cells in AAA. For T cells, we identified multiple clones in 11 of 16 AAA samples and in 1 of 8 control samples. Comparison of the immune receptor repertoires indicated that only few clones were shared between the individual AAA samples. The most frequently used V-genes in the TCR beta chain in AAA were TRBV3, TRBV19, and TRBV12-2+TRBV13-2.

**Conclusion:** In summary, we found no clonal expansion of TCRs or BCRs in elastase-induced AAA in mice. Our findings imply that a more precise characterization of TCR and BCR distribution requires a more extensive amount of T and B cells to prevent undersampling and to enable detection of potential rare clones. Using this current scSeq-based approach we did not identify clonal enrichment of T or B cells in experimental AAA.

## Introduction

Abdominal aortic aneurysm (AAA) is a cardiovascular disease defined as permanent dilation of the abdominal aorta greater than 50% or 3 cm [1]. Most AAAs develop in the infrarenal region between the renal veins and the aortic bifurcation [2]. The prevalence of AAA is 4-8% in men older than 60 years and 0.5-1.5% in women, with the rupture of AAA being associated with a high mortality rate [3]. AAA is a multifactorial and progressive disease. Genetic factors and inflammation strongly contribute to AAA development and in the last few years several studies have revealed that autoimmunity may contribute to the pathogenesis of AAA [4].

Inflammation and immune cell recruitment are characteristic of AAA. Accordingly, T and B cells are among the predominant infiltrating immune cells in human AAA tissue are [5]. Presence of these lymphocytes in AAA tissue was confirmed in several experimental mouse models of AAA [6–10]. In addition, B cell-bound immunoglobulins, such as IgM and IgG, have been localized in AAA tissue, which promote inflammation and tissue degradation [6,7,9,11]. The majority of lymphocytes infiltrating AAA tissue are T helper-2 (Th2) cells [12]. Th2-cells release cytokines such as interleukin- (IL-) 4, 5, 8 and 10, Fas ligand, Fas associated phosphatase-1, Interferon gamma and CD40 ligand that are associated with macrophage activation, regulation of vascular smooth muscle cell apoptosis and remodeling of the aortic wall [3, 12]. Animal studies showed that IL-4, IL-10 or CD4 deficiency in mice lead to increased AAA size in elastase-induced AAA suggesting a crucial role of Th2-cells in the pathogenesis of AAA [12]. Investigation of the TCR/antigen/human leukocyte antigens (HLA) complex revealed evidence that AAA may be a specific antigen-driven T cell disease [5, 13]. Studies discovered clonal expansion of T cell clones in AAA lesions, linked AAA to specific HLA Class I and Class II antigens, and identified self or nonself antigens that may be associated with AAA [5].

The role of B cells in AAA is still controversially discussed. Depletion of B cells can prevent AAA growth in different experimental aortic aneurysms. Shaheen et al. showed that depletion of B cells with an anti-CD20 antibody suppressed AAA growth in the angiotensinII- and the elastase perfusion AAA model [14]. Furusho et al. could corroborate the findings in B cell-deficient muMT mice, which underwent AAA induction by periaortic application of CaCl_2_ [7]. The application of polyclonal IgG antibodies to B cell-depleted mice resulted in larger AAA size comparable to wild-type (WT) mice indicating that IgG is sufficient for AAA development [7]. In contrast, Meher et al. could not observe differences in experimental AAA formation between B cell-deficient and WT mice and further showed that adoptive transfer of B2 cells suppressed AAA formation and decreased infiltration of mononuclear cells into aneurysmal tissue upon adoptive transfer of B2 cells [8]. However, there is evidence that an autoimmune process directed against self-antigens in the aortic wall may play a role in AAA pathogenesis. Zhou et al. identified a natural IgG antibody against fibrinogen in aortic tissues of elastase-induced AAA that induces AAA formation by activating the complement lectin pathway [15]. Further investigation of the role of B and T cells and the immunoglobulins involved in AAA is essential to gain a better understanding of the pathogenesis underlying AAA. Clonality and diversity of the immune receptor repertoire provide insights into disease mechanisms. They further can predict the immunological status of an individual and therefore be used for disease diagnosis [16].

Adaptive immune receptor repertoire (AIRR) sequencing is increasingly used to investigate lymphocyte dynamics in pathological contexts such as autoimmune and sterile inflammatory diseases, cancer and infections [17]. T cell receptors (TCR) and B cell receptors (BCR) are highly diverse heterodimers that recognize an immense variety of antigens [18]. The receptors consist of a combination of heavy and light chains in case of BCRs and a combination of α/β or γ/δ chains in case of TCRs. The majority of TCRs expressed on T cells consist of a combination of α and β chains. The receptors are formed by V(D)J recombination, which is the rearrangement of the variable (V), diversity (D) and joining (J) gene segments. For TCR α chains and BCR light chains, only V- and J-genes are involved in the recombination. Additional diversity is achieved by addition or deletion of random nucleotides at the junction sites between the gene segments and by the chain pairing. In case of BCRs, somatic hypermutation also results in greater diversity. Each receptor chain contains three hypervariable loops termed complementarity determining regions (CDR) that are required for interaction of the receptors with the antigen. CDR3 is commonly used as a region of interest to determine T and B cell clonotypes due to its high diversity and essential role for antigen binding. [18-21] A clonotype is a set of cells expressing the same immune receptor, which implies that the receptors consist of the same V- and J-genes and encode an identical CDR3 amino acid sequence [22]. The AIRR is the union of all TCRs and BCRs of one individual and can change greatly with onset and progression of diseases [18]. The TCR repertoire within one individual is estimated at 10^7^ TCRs in humans and 10^6^ TCRs in mice [23] while the estimated size of the B cell repertoire is 10^18^ in humans and 10^13^ in mice [24, 25].

Previously, TCRs were analyzed in aortic aneurysms and their function has been investigated in mice and humans, yet no study addressed B cell clonality in aortic aneurysms. Li et al. found clonal expansion of regulatory T cells (Treg) in mouse aortae after elastase-induced AAA formation [26] and several studies showed the presence of clonally expanded TCRs in aneurysmal lesions of patients with AAA or ascending thoracic aortic aneurysms, supporting the notion that AAA may be promoted by specific-antigen driven T cells [13,27–29].

Single-cell RNA (scRNA) sequencing of TCRs and BCRs is a powerful tool to investigate the immune receptor repertoire involved in AAA pathology. In comparison to bulk RNA sequencing, where a mixture of different gene expression profiles is obtained from the material studied, scRNA sequencing offers several advantages [30]. scRNA sequencing provides information of TCR chain pairing, higher resolution, and is more suitable for investigating the TCR specificity for an antigen of interest [31]. However, there are also some limitations of scRNA sequencing. These challenges include the isolation of living single cells out of tissues, the lower output of sequenced cells compared to bulk sequencing and higher costs [31]. In particular, for RNA sequencing of human AAA only a small number of cells of interest is available for analysis, as only small sections of AAA can be collected during surgery [32]. In mouse models, the whole AAA can be used, but the total amount of T and B cells is still small for scRNA TCR and BCR sequencing. Zhao et al. obtained approximately 3000 cells, encompassing all present cell types, from topical elastase-induced AAA of 10 mice [33]. Due to the small number of cells, biological interpretation of the sequenced TCR and BCR repertoire in AAA is challenging. In addition, standardized and uniform sample preparation, pre-processing of data and bioinformatics workflow for data analysis is important to obtain robust and comparable data. There are already several guides [34], tools [35], and pipelines [36] for data analysis. In this paper, we highlight problems of scRNA TCR and BCR sequencing specifically in AAA and provide a strategy for performing these experiments as well as for subsequent data analysis using a dataset we generated as an example.

## Methods

### Mice

Male C57BL/6J mice purchased from Janvier Labs (Saint-Berthevin, France) were used for experiments at the age of 10-11 weeks. All animal procedures were performed in accordance with the European convention for the protection of vertebrate animals used for experimental and other scientific purposes and were approved by LANUV (Landesamt für Natur, Umwelt-und Verbraucherschutz Nordrhein-Westfalen; AZ 81-02.04.2018.A408). Mice were housed under standard laboratory conditions with a 12 h light/dark cycle and had ad libitum access to drinking water and standard chow.

### Porcine pancreatic elastase perfusion model

To induce AAA in mice the porcine pancreatic elastase (PPE) perfusion model was used as previously described by Pyo et al. [37]. In brief, mice received analgesic by injecting 0.1 mg/kg body weight (bw) buprenorphine subcutaneously prior to surgery. Mice were anesthetized with isoflurane (initial 3%, then 1.5%) and oxygenated air. After absence of the toe reflex, laparotomy was performed, the proximal and distal infrarenal aorta was isolated, and temporarily ligated. The aorta was punctured, a catheter was inserted and the infrarenal part was perfused with sterile isotonic saline containing type I porcine pancreatic elastase (2.5-3 U/ml Sigma Aldrich, #E1250) or 0.9% NaCl (sham surgery) under 120 mmHG for 5 min. Elastase concentrations ranged from 2.5 to 3 U/ml depending on the batch number, as different concentrations were necessary to trigger the same AAA incidence and size. The aortic puncture was sutured, the ligations were removed and the abdomen was closed. Afterwards the mice received buprenorphine (0.1 mg/kg bw, subcutaneously) if required in the first eight hours. Additionally, mice received buprenorphine (0.01 mg/ml) via the drinking water for three days. Mice were monitored regularly until the end of the experiment.

### Organ harvesting

Infrarenal aortae were harvested on day 3 (n = 5), 7 (n =5), 14 (n = 2) and 28 (n = 4) after PPE surgery and on day 3 (n = 3) and 28 (n = 3) after sham surgery. Additionally, for two control samples infrarenal aortae from three non-treated C57BL/6J mice were pooled. In total we obtained 24 samples for scRNA sequencing. Approximately 10 min prior organ harvesting mice were injected i.v. with 100 µl CD45-FITC antibody (#553079, Biolegend, dilution 1:1000) to label circulating leukocytes. Mice were anesthetized and analgesized with ketamine (100 mg/kg bw) and xylazine (10 mg/kg bw). After absence of the toe reflex blood was collected from the heart with a heparinized syringe. Thorax and abdomen were opened, the vena cava was cut and the cardiovascular system was perfused with cold PBS through the left ventricle of the heart. The infrarenal part of the aorta was isolated by carefully removing all fatty tissue, collected and stored in PBS on ice until further processing.

### Digestion of aortic tissue into single cells

The isolated infrarenal aortae were digested into single cells based on the protocol from Hu et al. [38]. Briefly, aortae were cut and transferred into an enzyme mix containing 400 U/ml Collagenase I (Sigma Aldrich, #C0130-100MG), 120 U/ml Collagenase XI (Sigma Aldrich, #C7657-25MG), 60 U/ml Hyaluronidase I-S (Sigma Aldrich, #H3506-100MG) and 60 U/ml Dnase I (Sigma Aldrich, #46944200) in Dulbecco’s phosphate buffered saline (DPBS) containing calcium and magnesium supplemented with 20 mM HEPES (Thermo Fisher Scientific, #15630106). Aortae were incubated in the enzyme mix for 50 min on a shaker (600 rpm) at 37 °C. The cell suspension was filtered through a 100 µm cell strainer (pluriSelect Life Science, Leipzig). The remaining aortic tissue was mashed with a syringe plunger through the cell strainer, which was rinsed several times with DPBS. After centrifugation (10 min, 450 x g, 4 °C) cells were resuspended in cold PBS and transferred into a 96-well plate.

For additional flow cytometer analysis only cells were subsequently resuspended in RPMI 1640 (Sigma Aldrich) supplemented with 10% fetal calf serum (Sigma Aldrich) and incubated on a shaker (600 rpm, 12 min, 37 °C). Finally, cells were centrifuged (10 min, 450 x g, 4 °C), resuspended in PBS and transferred into a 96-well plate.

### Staining of single cells

The 96-well plate was centrifuged for 5 min at 500 x g and 4 °C. Cells were stained with a staining mix containing Fc receptor blocker (TruStain fcX™, BioLegend, Amsterdam, The Netherlands, 1:100), viability stain (Zombie Aqua™ and Zombie Green™ Fixable Viability Kit, Biolegend, 1:500), CD45-AP/Cyanine7 (Biolegend, clone 30-F11, dilution 1:200), TER-119-FITC (Biolegend, clone TER-119, 1:200) and C0443 CD41 (Biolegend, Barcode Sequence ACTTGGATGGACACT, 1:1400) in PBS. Additionally, an individual TotalSeq Hashtag antibody (Biolegend, TotalSeq™-C) was added to the single-cell suspension of each mouse, respectively. The hashtag antibodies allow to combine samples from several mice in the same 10X sequencing run and are needed to demultiplex cells from individual mice. In order to have a hashtag for each mouse, 10 mice were hashtagged with an antibody from the TotalSeq™-C series (Biolegend) and two hashtag antibodies were built by combining the antibodies MHC I-Biotin (Biolegend, clone 28-8-6) and CD45-Biotin (Biolegend, clone 30-F11) with either TotalSeq™ C971 or C972. The samples were stained with the staining mix and hashtag antibodies for 15 min at room temperature (RT) in the dark. After centrifugation (5 min, 500 x g, 4 °C) supernatant was discarded and cells were resuspended in MACS buffer (Miltenyi Biotec, #130-091-221) for following cell sorting.

Only for flow cytometer analysis, CountBright™ Absolute Counting Beads (Thermofisher Scientific) were added to every sample prior to staining to determine cell counts. Cells were centrifuged for 5 min at 500 x g and 4 °C and stained with Fc receptor blocker (TruStain fcX™, BioLegend, 1:100) and viability stain (Zombie Aqua™ Fixable Viability Kit, Biolegend, 1:500) at RT for 10 min in the dark. After centrifugation (5 min, 500 x g, 4 °C) cells were stained with the following conjugated antibodies for 20 min at RT in the dark: CD3-APC (Biolegend, clone REA613, 1:200), CD11b-APC-Cy7 (Biolegend, clone M1/70, 1:200), CD19-FITC (Biolegend, clone MB19-1, 1:100), CD45-V450 (Biolegend, clone 30-F11, 1:200), Ly6C-PE (Biolegend, clone HK1.4, 1:133), Ly6G-PerCP-Cy5 (Biolegend, clone 1A8, 1:100), NK1.1-PE-Cy7 (Biolegend, clone PK136, 1:200). Cells were centrifuged (5 min, 500 x g, 4 °C) and resuspended in PBS with 0.5% bovine serum albumin. Samples were acquired with the BD FACSVerse™ Cell Analyzer (BD, Heidelberg, Germany) and data was analyzed with the FlowJo software v10.5.3.

### Cell sorting

Cell sorting was performed on a MoFlo XDP (Beckman Coulter, Krefeld, Germany). Every sample was sorted separately. For every sample 3,000 living CD45^+^ cells were sorted. If samples did not reach the appropriate cell count, this was compensated by sorting more cells of other samples. Cells from all samples were combined into one reaction tube and centrifuged for 5 min at 500 x g at RT. The supernatant was removed and cells were resuspended in MACS buffer. Recounting of the cells revealed approximately 60,000 living cells.

### Generation of single cell library

Single cell libraries were generated on the 10X Chromium Controller system utilizing the Chromium Next GEM Single Cell 5’ Kit v2 (10X Genomics, Pleasanton, CA, USA) according to manufacturer’s instructions. Sequencing was carried out on a NextSeq 550 system (Illumina Inc. San Diego, USA) with a mean sequencing depth of ∼50,000 reads/cell for gene expression. The T cell and B cell libraries as well as the hashtag libraries were sequenced at ∼5,000 reads/cells.

### Processing of 10X Genomics single cell data

Raw sequencing data was processed using the 10X Genomics CellRanger software (v6.0.2). Raw BCL-files were demultiplexed and processed to Fastq-files using the CellRanger *mkfastq* pipeline. Alignment of reads to the mm10 genome as well as the corresponding V(D)J gene references and UMI counting was performed via the CellRanger *multi* pipeline to generate a gene-barcode matrix.

### scRNA sequencing data analysis

R package *Seurat v4*.*0* [39] was used for analysis. First the hashtag library was added as assay to the metadata of the RNA library. Cells with less than 200 RNA counts and more than 30% mitochondrial RNA were excluded. Data was normalized and scaled with the functions *NormalizeData(), ScaleData()*. Principal component analysis, variable genes finding, cell clustering and Uniform Manifold Approximation and Projection (UMAP) dimensional reduction was performed. Doublets were removed with positivity for more than one hashtag and *DoubletFinder v2*.*0* [40]. T cells were defined by expression of Cd3e, Cd3d, Cd3g, Cd28. B cells were defined by expression of Cd19, Cd79a, Cd79b. T and B cells were isolated bioinformatically and merged with the preprocessed scRNA TCR and BCR sequencing data.

### scRNA TCR and BCR sequencing analysis

Preprocessing of the scRNA TCR and BCR sequencing data included quality control and adding the library of the hashtag antibodies. Only receptors with exactly one alpha/heavy chain and one beta/light chain were used for analysis. Sequences of single chains, more than two chains per receptor and not matching chains were excluded from the analysis. Then, the hashtag information was added to the 10x output file, to enable assignment of the TCRs and BCRs to the different mice. The *Immunarch package v 0*.*6*.*9* [41] was used to analyze CDR3 length distribution, clonotype abundance, repertoire overlap, germline gene V-gene usage, clonal expansion and diversity estimation.

### Statistics

Data are presented as absolute numbers or mean ± s.d. Two-sample permutation-based Kolmogorov-Smirnov test was used to compare distributions with the function ks_test from R package “twosamples”. One-sided Fisher’s exact test with Bonferroni correction was used for database comparisons with R v4.0 software. Results with p ≤ 0.05 were considered significant.

## Results

### scRNA sequencing Workflow

The overall goal of our scRNA sequencing experiments was to decipher lymphocyte dynamics and heterogeneity in elastase-induced aneurysms in mice at different disease stages and to additionally investigate TCR and BCR clonality. This paper focuses on the results of TCR and BCR sequencing, highlights problems and pitfalls of these types of experiments, and provides guidance for future studies.

The experimental workflow starts with the induction of AAA by perfusing the infrarenal aorta of male C57BL/6J mice with elastase or NaCl (Figure 1A). Infrarenal aortae were harvested on day 3 (n = 5), 7 (n =5), 14 (n = 2) and 28 (n = 4) after PPE surgery and on day 3 (n = 3) and 28 (n = 3) after sham surgery to cover different disease stages. Additionally, infrarenal aortae from 6 non-treated C57BL/6J mice of which three were pooled into one sample were harvested for sequencing. In total 24 samples were subjected to scRNA sequencing and immunoreceptor analysis. Organ harvesting, digestion of aortic tissue into single cell suspensions as well as staining and sorting of single cells was performed as described in the method section. Then, scRNA sequencing and TCR and BCR scRNA sequencing was performed.

**Figure 1:**
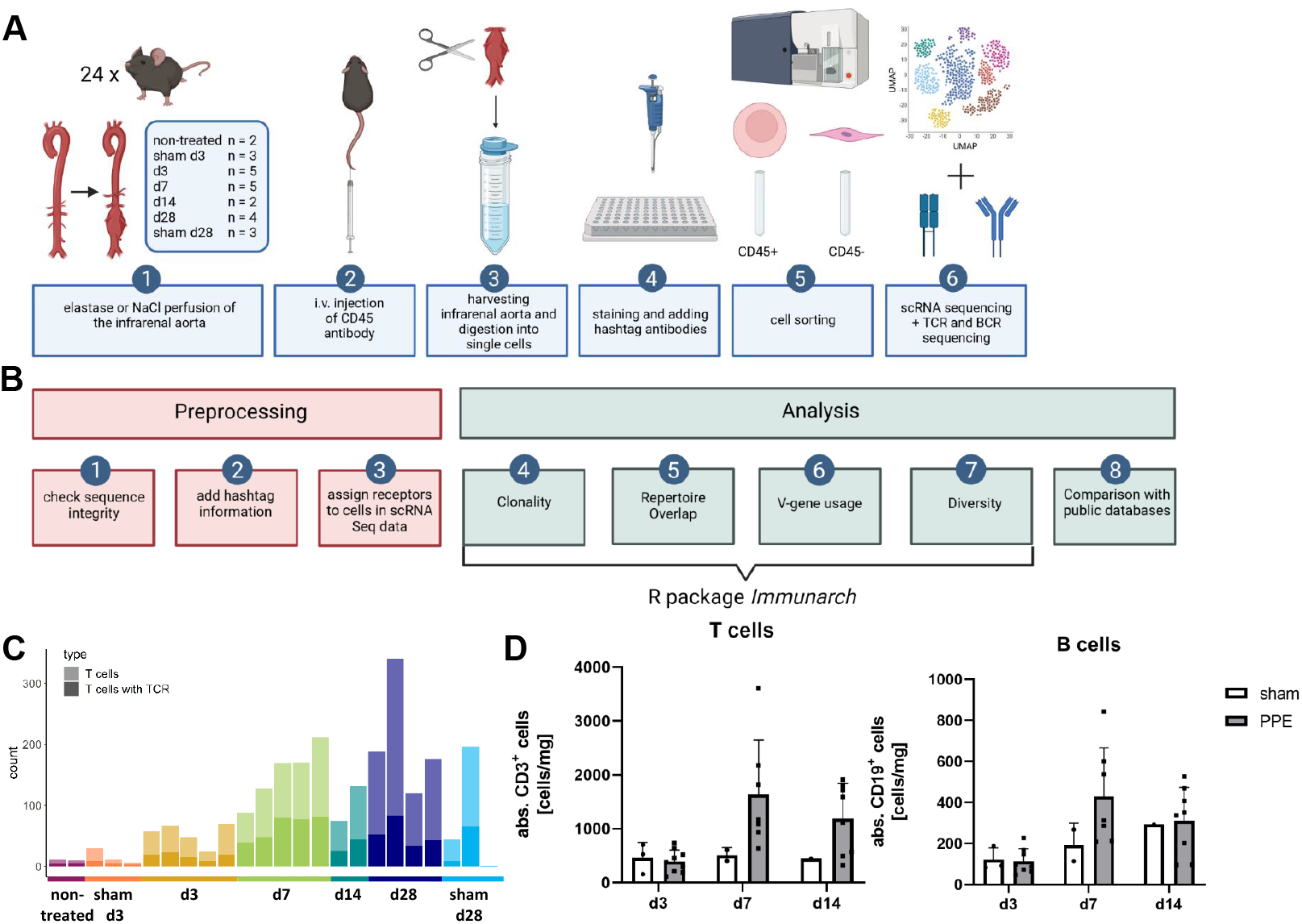
Experimental and bioinformatic workflow as well as quality control of the data. **A:** Experimental workflow. AAA was induced in mice via perfusion of the infrarenal aorta with elastase. NaCl perfusion of the aorta served as sham-operated control. Prior to organ harvesting mice were intravenously injected with a fluorophore labeled CD45 antibody. Harvested infrarenal aortae were digested into single cells and stained with antibodies and hashtags. Cells were sorted for living leukocytes and living non-leukocytes. scRNA sequencing as well as TCR and BCR scRNA sequencing was performed. **B:** Bioinformatic workflow including preprocessing and data analysis. Three preprocessing steps were performed as quality control for the data. Only fully intact sequences are retained for analysis. The information for the hashtag antibodies was added to the TCR and BCR scRNA sequencing data. The receptors were assigned to the corresponding cells in the scRNA sequencing data set. The main data analysis including clonality, repertoire overlap, V-gene usage and diversity was performed with the *immunarch* R package. In addition, the data was compared with public databases. **C:** T cell amounts (light bar color) and T cell amount exhibiting an intact TCR (dark bar color) in aortic tissue on different AAA disease stages received from scRNA sequencing data. The amount of T cells increases with AAA progression. **D:** Flow cytometer analysis of T and B cell amounts of aortic tissue 3,7 and 14 days after sham operation or PPE-induced AAA formation.

Consistent preprocessing of immune receptor scRNA sequencing data is crucial for comparable data. We suggest using only those immune receptor data, with all fragments intact and both chains (a/ß for TCR or heavy/light for BCR) present. Receptors with only one, or more than two chains, that can appear in the data due to sequencing errors, were excluded from the data. After that, the information of the hashtag antibodies was added to the data to assing an immune receptor specifically to one cell of a specific mouse. Subsequently, the receptors were assigned to the corresponding cell in our scRNA sequencing dataset. The majority of the immune receptor analysis such as clonality, repertoire overlap, V-gene usage and diversity was performed with the R package *immunarch* [41]. In addition, comparison of the data with public databases to identify disease associated receptors was performed (Figure 1B).

### Analysis of scRNA TCR and BCR sequencing data

To ascertain quality control, we processed the TCR and BCR sequencing data and evaluated basic statistics. The raw data contained 3370 TCR sequences (1484 TCR alpha chains (TRA), 1886 TCR beta chains (TRB)) and 1745 BCR sequences (570 immunoglobulin heavy chains (IGH), 1131 immunoglobulin kappa (IGK) light chains, 44 immunoglobulin lambda (IGL) light chains. Checking the integrity of the immune receptors yielded a high number of immune receptors with only one sequenced chain and some receptors with more than two chains, which were excluded (Supplement Figure 1). After that step 2296 TCR chains (1148 pairs of TRA and TRB chains) and 980 BCR chains (490 pairs of heavy and light chains) remained. We next filtered for receptors that could be associated with a hashtagged cell and retained 2012 TCR chains and 770 BCR chains. Assignment of the immune receptors to the corresponding cells in our scRNA sequencing data revealed that only 46.49 % of BCRs were expressed in B cells (defined by expression of Cd19, Cd79a, Cd79b), and the majority of TCRs (79.13 %) was expressed in T cells (defined by expression of Cd3e, Cd3d, Cd3g, Cd28), whereas the remaining immune receptors are found on other cell types (Supplement Figure 1). TCRs and BCRs not expressed in the respective lineage were excluded to avoid analysis of false positive receptors due to sequencing artifacts. The final analysis included 1592 TCR chains (796 pairs of TCRs) and 358 BCR chains (179 pairs of BCRs).

We next compared the number of immune receptors with that of T and B cells that are present in our scRNA sequencing dataset and displayed the distribution across the timepoints and samples (Figure 1 C, Supplement Figure 2). Overall, there were less B cells (325) than T cells (2376) in AAA tissue and 3 out of 24 samples did not contain B cells (non-treated, d3, sham d28) (Supplement Figure 2). The amount of both cell types increased with advanced disease stages and highest levels were detected at d28 (Figure 1 C and Supplement Figure 2). We corroborated our findings by flow cytometry revealing a time- and disease stage-dependent increase of lymphocytes in AAA. Of note, only few cells were detected in sham operated mice (Figure 1 D). A TCR could be assigned to 33.5% of the present T cells (2376 T cells, 796 TCRs). Thus, a large proportion of TCR sequences present in AAA are missing due to inefficient sequencing. In our data 55.08% of B cells have a matching BCR (325 B cells, 179 BCR). In 4 out of 24 samples no BCRs could be detected, these were sham-operated or early timepoints (non-treated, sham d3, d3, sham d28) (Supplement Figure 2).

### Estimating clonal expansion by spectratyping

Spectratyping identifies the pattern of the CDR3 length distribution [42]. Comparing the shape of the CDR3 length distribution between control and disease can indicate presence of clonal expansion in a repertoire. Deviations from the normal pattern might be due to the high frequency of a specific CDR3 sequence and are therefore associated with clonal expansion [20]. We compared the CDR3 length distribution of TCRs in AAA, including all timepoints, and controls, including sham-operated and non-treated mice, using two-sample permutation-based Kolmogorov-Smirnov test (Figure 2A). The resulting p-value of 0.92 suggests that the CDR3 length follows the same distribution in AAA and controls. We performed the same analysis to compare the CDR3 length distribution between the different disease stages yielding also no significant differences (Figure 2B).

**Figure 2:**
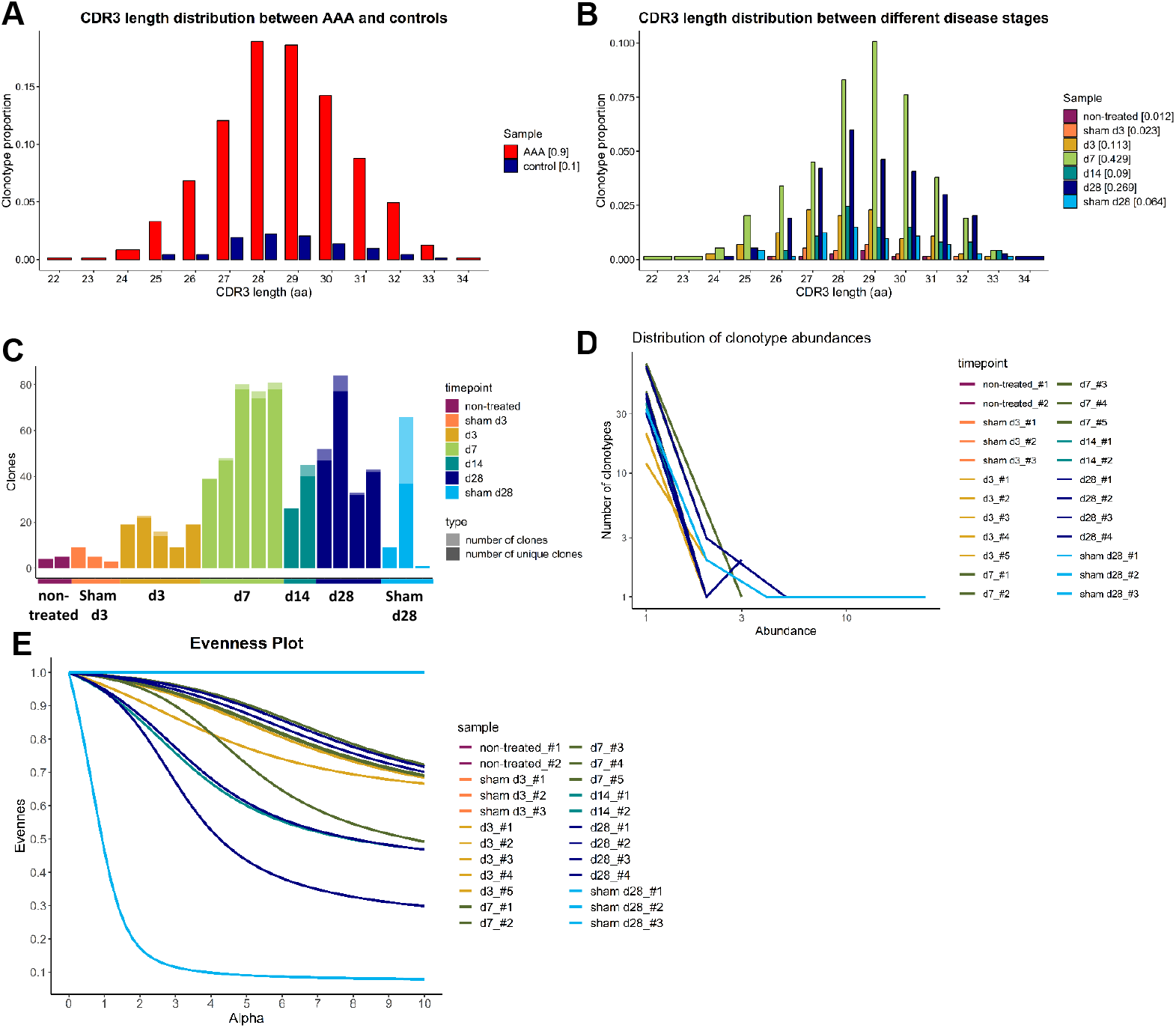
CDR3 length distribution and clonotype abundance reveals no clonal expansion of TCRs in elastase-induced aneurysm in mice. **A:** No changes in CDR3 length distribution of TCRs (paired chains) between AAA including all different disease stages (red) and control samples including sham operated and non-treated mice (blue). The clonotype proportion is plotted against the amino acid (aa) CDR3 length. **B:** No alterations in CDR3 length distribution of TCRs (paired chains) between samples of different disease stages (d3, d7, d14, d28) and sham operated as well as non-treated samples (different colors). The clonotype proportion is plotted against the amino acid (aa) CDR3 length. **C:** Amount of all TCR clonotypes per sample (light bar color) including the amount of unique clones (dark bar color). The majority of TCR clones were found to be unique. Multiple copies of one clone appear only in one of the sham_d28 samples and in 11 the AAA samples. **D:** Line Plot indicating the number and abundance of clonotypes per sample. In the sham_d28 sample exhibiting clones with multiple copies, one clone is present 30 times. Whereas, in AAA samples one clone only appears 2 to 6 times. Abbreviations: CDR = complementarity determining regions. **E**: Evenness Plot indicating the extent of clonal expansion for every sample.

Receptor clonality can also be investigated by determining the number of unique clones and clonotype abundance. A clonotype was defined as receptors that consist of the same V- and J-genes, and encoding an identical CDR3 amino acid sequence. The majority of TCR clones in AAA, sham-operated and non-treated aortae are unique (Figure 2C). Only one sample of the sham operated and non-treated aortae and 11 AAA samples exhibit expanded clonotypes (Figure 2C). However, the same clonotype is infrequent in AAA samples and occurs 2 to 6 times, whereas the one sham-operated sample exhibiting several clonotypes, has one T cell encompassing 30 cells (Figure 2D).

The extent of receptor clonality can be indicated with an evenness profile of the repertoire [16]. The alpha values represent different diversity indices with different weights on expanded clonotypes. Higher alpha values give more weight to expanded clonotypes, while alpha = 0 weights every clonotype equally regardless of its frequency. Therefore, high receptor clonality is indicated as a highly uneven curve and no receptor clonality is associated with a completely even curve. The sham operated d28 sample, in which one clonotype was identified 30 times (Figure 2D), exhibits also the highest clonality, whereas all other sham operated or non-treated samples show no clonality (Figure 2E). Additionally, 11 AAA samples from different timepoints show a lower extent of receptor clonality. The evenness profiles of the individual samples are displayed in Supplement Figure 3.

### Investigating the TCR repertoire similarity

Repertoire overlap analysis is commonly used to identify “public” clonotypes that are shared between individuals. The R package *immunarch* provides several methods to measure receptor similarities between individuals. Using the function “public” specified the exact number of shared immune receptors between different repertoires, thereby revealing that in 7 instances a TCR sequence is equal in 2 AAA samples (Figure 3A). Additionally, repertoire similarity can be investigated by identifying TCRs of different individuals containing the same V-region genes. Fragments of V-region genes are classified into families according to their nucleotide sequence similarity (at least ∼70%). Specific V-gene usage patterns have been associated with different diseases and were shown to change in response to therapeutic approaches [43, 44]. In our data the V-gene usage of the beta chain (TRBV) correlates stronger than the V-gene usage of the alpha chain (TRAV) of the TCR (Figure 3B). A deeper analysis of the distribution and frequency of used TRBV genes revealed a high usage of TRBV3, TRBV19, and TRBV12-2+TRBV13-2 in AAA samples at day 7, 14 and 28 (Figure 3C).

**Figure 3:**
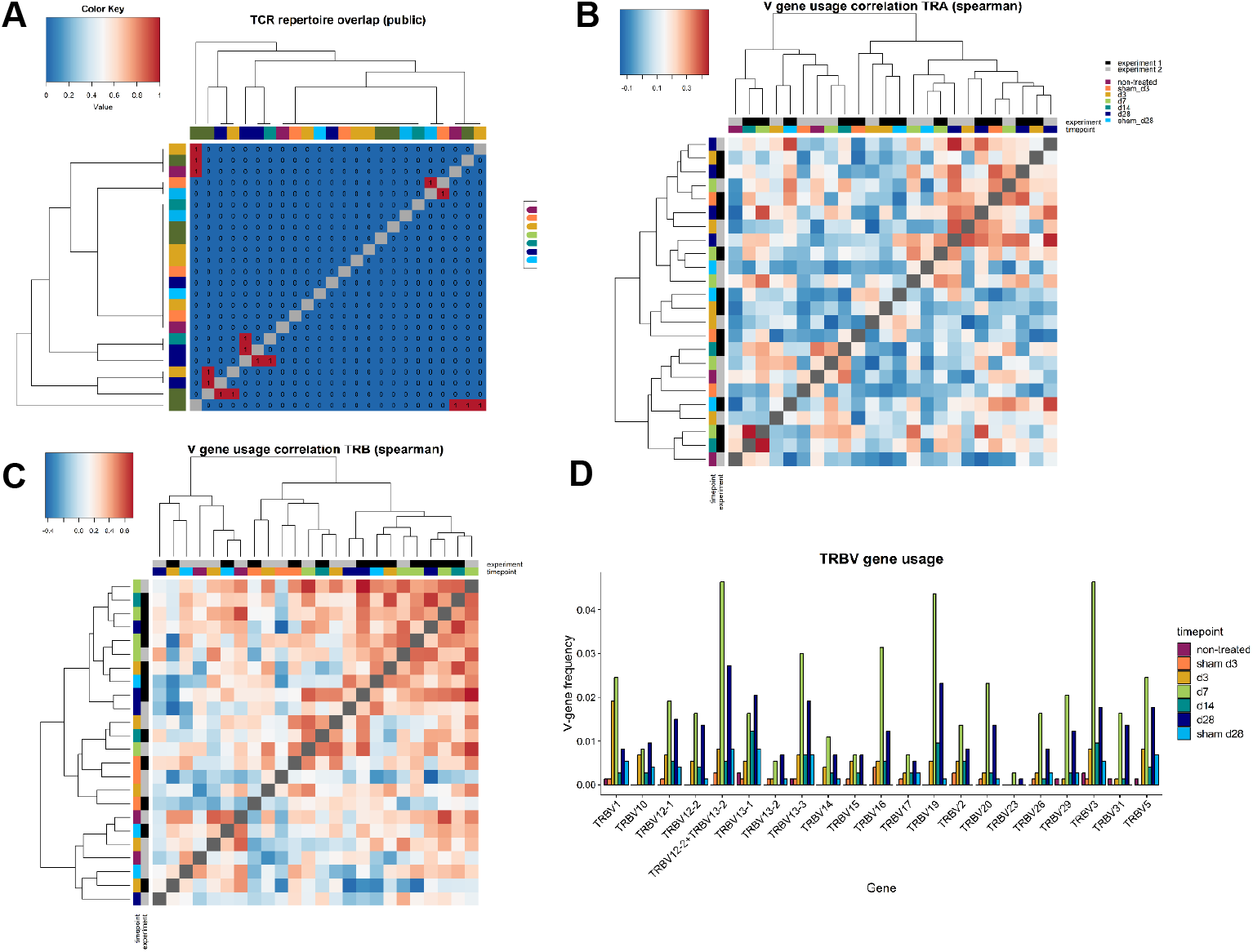
Comparing the different TCR repertoires reveals shared TCR sequences, a high correlation of the V-gene usage and several frequently used TRBV genes in AAA. **A:** Analysis of the TCR repertoire overlap shows that in 7 instances two AAA samples contain one equal TCR. The color code in the heatmap indicates the different samples. **B:** The V-gene usage of the TCR alpha chain (left heatmap) and of the TCR beta chain (right heatmap) is correlated and hierarchical clustered between the different samples using spearman correlations. Color gradient indicates the level of correlation (blue = negative correlation, red = positive correlation). The color code on the axes indicates the different samples. **C:** Distribution and frequency of TRBV genes occurring in all samples. Frequently used TRBV genes in AAA samples are TRBV3, TRBV19, and TRBV12-2+TRBV13-2 at day 7, 14 and 28. Abbreviations: TRA = TCR alpha chain ; TRAV = TCR alpha chain v gene; TRB = TCR beta chain; TRBV = TCR beta chain v gene

Frequently used TRBV genes in AAA samples are TRBV3, TRBV19, and TRBV12-2+TRBV13-2 at day 7, 14 and 28. Abbreviations: TRA = TCR alpha chain ; TRAV = TCR alpha chain v gene; TRB = TCR beta chain; TRBV = TCR beta chain v gene.

### Dataset comparison with public TCR and BCR databases

We compared the presence of CDR3 sequences in our dataset with the two public TCR databases VDJdb [45] and McPAS-TCR [46] to investigate if TCR clones in our dataset are associated with other diseases or antigens [47]. VDJdb is a curated database of TCR sequences with known antigen specificities containing TCR information of different species (*Homo Sapiens, Macaca Mulatta* and *Mus Musculus*) and various diseases [45]. McPAS-TCR is a database of TCR sequences found in T cells that were associated with various pathological conditions in humans and in mice [46]. TCRs with less than 4 amino acids and diseases with less than 5 TCRs were excluded. Furthermore, categories within the MusMusculus databases, which were not relevant for comparison, like “Synthetic” or “GallusGallus” were excluded. Accordingly, for VDJdb 5206 TCRs found in Influenza (3156 TCRs), Lymphocytic choriomeningitis virus (LCMV) (151 TCRs), Murine cytomegalovirus (MCMV) (1463 TCRs), Plasmodium Berghei (245 TCRs), Respiratory syncytial virus (RSV) (125 TCRs) and Vesicular stomatitis virus (VSV) (66 TCRs) and for McPAS-TCR, 3530 TCRs which are assigned to 21 different diseases/pathogens were used for analysis (Supplement Table 1). After merging the two databases and filtering for unique CDR3 sequences, we obtained 4331 CDR3 sequences for comparison with our data set. One-sided Fisher’s exact test was used to examine the overrepresentation of TCR clones in our data set that are associated with diseases or antigens according to the two databases. Our dataset shared 56 CDR3 sequences with the public databases which were assigned to MCMV (p = 0.006), LCMV (p = 0.01), Influenza, Plasmodium Berghei, VSV, Diabetes type 1 and Tumor (Figure 4B and Supplement Table 2). The obtained p-values were adjusted for multiple testing using Bonferroni correction. Bonferroni correction resulted in no significant p-values indicating there are no TCR clones overrepresented in our data set that are associated with diseases or antigens according to the two databases.

**Figure 4:**
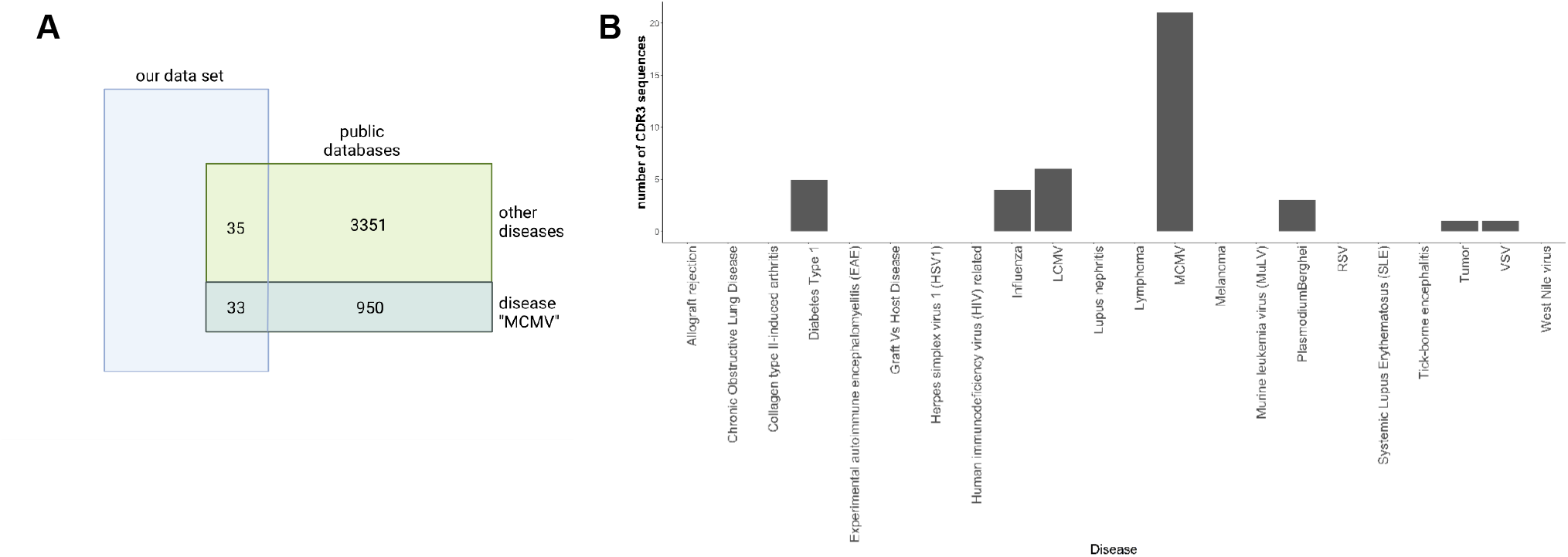
Overlap of AAA-associated CDR3 sequences with public databases. **A:** Venn diagram indicating the overlap of our data set with the database representative for the disease MCMV. Our data shares 33 CDR3 sequences with the public databases that are associated with MCMV and 35 CDR3 sequences that are associated with other disease. **B:** Barplot displaying the amount of sequences per disease in our dataset

### No clonal expansion or repertoire overlap of BCRs in elastase-induced aneurysm in mice

The CDR3 length distribution of the BCRs shows no differences between AAA and control at the disease stages (Figure 5 A). Most BCR clones are unique. As mentioned before, BCRs are present in only 20 of 24 samples in our dataset. The samples that lack BCRs are control samples or early disease stages that are known to contain few B cells overall (non-treated, sham d3, sham d28, d3). Clonotypes that appear more than once were found in two of these 20 samples. One of the day 28 samples contains one BCR that is present thrice, the day 7 sample has one BCR that appears twice (Figure 5 B). The evenness profile likewise indicates clonal expansion for these two AAA samples, whereas all other samples show no clonality (Figure 5C). Next, we investigated the isotype distribution of the BCR heavy chains. The most frequent isotype is IGHM, followed by IGHD (Figure 1D). The similarity measurement of the BCR repertoire present in the different samples shows that two AAA samples (d7 and d28) share 1 BCR (Figure 1E). Otherwise, there is no similarity between the different samples. The correlation of V-gene usage between the different samples is likewise low (Figure 1F). The highest correlation is found between one non-treated control and one d3 sample (r = 0.701). Overall, these data reveal no evidence for clonality among B cells in AAA.

**Figure 5:**
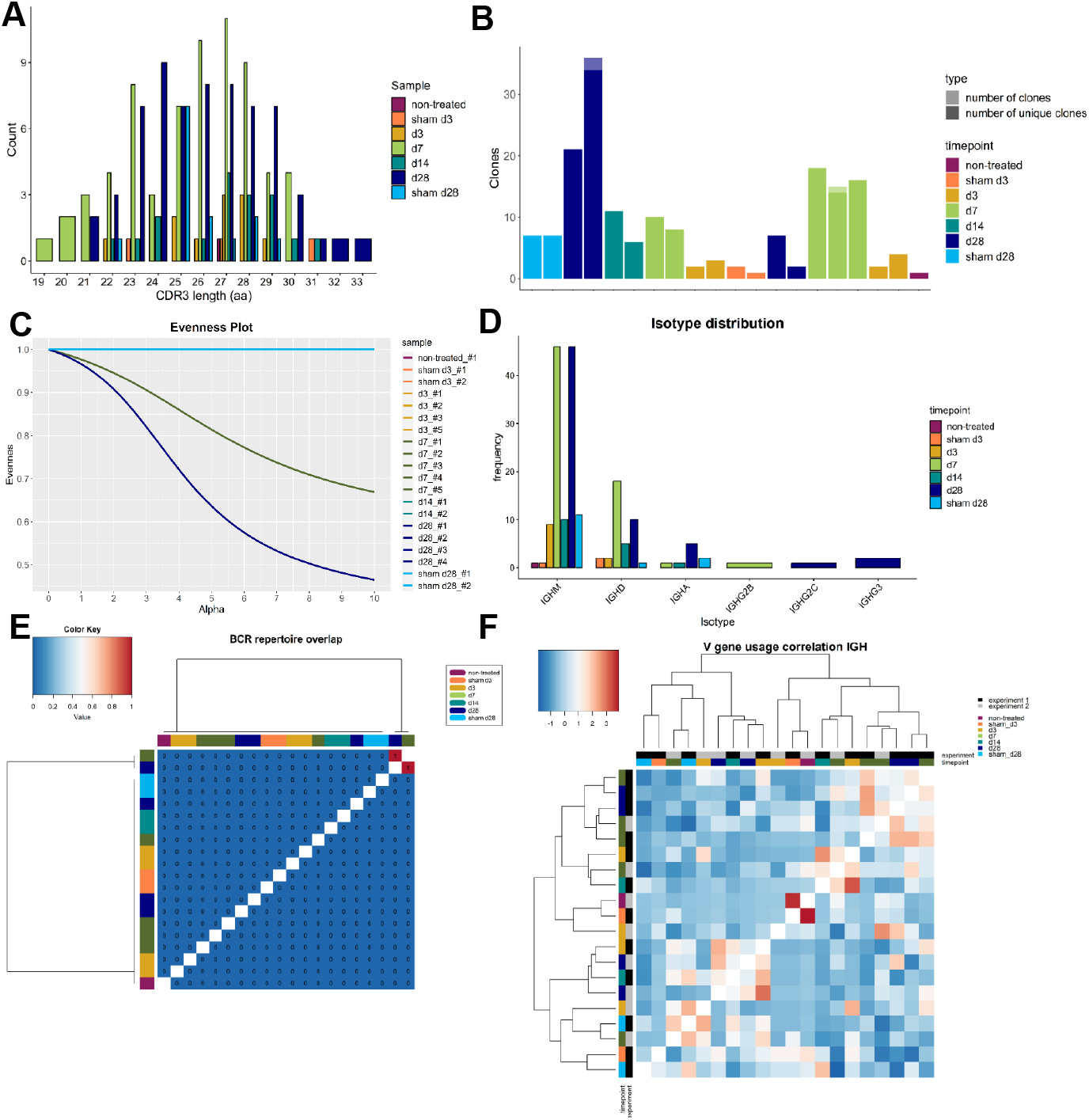
Elastase-induced AAA shows no BCR clonality. **A:** Histogram indicating the CDR3 length distribution of the BCRs (paired chains). The clonotype proportion is plotted against the amino acid (aa) CDR3 length. The color indicates whether the TCRs belong to an aorta isolated at d3, d7, d14 or d28 after PPE or sham surgery or to a non-treated aorta. **B:** Barplot displaying the number of all BCR clones (pale color) and only unique clones (vibrant color) per mice. Almost all BCR clones are unique. **C:** Evenness Plot indicating the extent of clonal expansion for every sample. **D:** Barplot indicating the isotype distribution of the BCR heavy chain (IGH) in the different conditions. Most frequent isotypes for all conditions are IGM and IGD. **E:** BCR repertoire overlap displayed in a heatmap. Color bars on the top and left side of the heatmap indicate the time point. **F:** Heatmap presenting the spearman correlation and hierarchical clustering of the BCR heavy chain (IGH) V-gene usage of the different samples. Color gradient indicates the level of correlation (blue = weak correlation, red = strong correlation). Color bars on the top and left side of the heatmap indicate the experiment and the time point.

## Discussion

We investigated TCR and BCR clonality in elastase-induced AAA in mice at different disease stages using scRNA TCR and BCR sequencing. Our results show no differences in CDR3 length distribution of TCRs and BCRs between the different disease stages, indicating no clonal expansion of immune cell receptors in elastase-induced AAA. The clonotype abundance likewise revealed no clonal expansion of BCRs in AAA. For the TCRs we found multiple clones in 68% of AAA samples and no clonality in control samples except for one. Comparison of the immune receptor repertoires showed a low similarity between the individual samples. Spearman correlation to compare the V-gene usage between the different AAA samples and controls revealed that the V-gene usage of the TCR beta chain correlates stronger than the V-gene usage of the TCR alpha chain. The most frequently used V-genes in the TCR beta chain in AAA are TRBV3, TRBV19, and TRBV12-2+TRBV13-2. Comparison of our TCR clones revealed no overrepresentation of TCR clones associated with diseases or antigens annotated in the 2 public databases. The main isotype in our BCR data is IgM followed by IgD. Both isotypes are expressed by naive B cells.

When immune cell receptors of naive lymphocytes recognize an antigen, the cell is activated and starts to proliferate. This process is termed clonal expansion and enables a targeted, adaptive immune response. Yet, the term or definition of clonal expansion still remains heavily debated. Lu et al. investigating T cell clonality in aneurysmal lesions of AAA patients, defined clonal expansion as presence of multiple identical copies of TCR transcripts. They reasoned that the size of the T cell repertoire makes it unlikely that multiple identical copies of a TCR transcript would be found by chance in an independent sample of T cells [27]. However, it is difficult to distinguish antigen-related clonotype expansion from convergent recombinations. Convergent recombinations arise due to biases in the VDJ recombination process during which some sequences are more likely to be generated than others. Thus, identical sequences appear multiple times in almost every individual and same sequences are also shared between different individuals [48, 49]. Furthermore, different combinations of inserted and deleted nucleotides can produce exactly the same sequence. Therefore, the occurrence of the few identical receptor sequences and the shared sequences between samples in our data set is likely due to convergent recombination rather than clonal expansion in response to a common antigen. In conclusion, we found no evidence for clonal expansion of TCRs or BCRs in elastase-induced AAA in mice. It must be considered that in this particular mouse model, aneurysm formation is triggered by enzymatic destruction of elastin fibers and does not fully mimic the complexity of AAA development in humans. In addition, our dataset contains only a few TCRs and BCRs, representing only a small portion of the actual repertoire in AAA, thus, making it difficult to draw an absolutely firm conclusion.

Li et al. for example induced AAA in mice with elastase and CaPO_4_, performed scRNA sequencing combined with TCR sequencing on 41341 CD4+ T cells and found a clonal expansion of regulatory T cells present in AAA [26]. Receptor diversity and abundance are correctly reflected in a sample only if the number of cells is sufficient [50]. This suggests that the number of analyzed cells is an important factor for investigating TCR clonality and a high number of cells facilitates the identification of clonal expansion.

Studies with patients demonstrated the presence of clonally expanded TCRs in AAA or ascending thoracic aortic aneurysms (TAA), supporting the notion that AAA may be an specific antigen-driven T cell disease [13, 27-29]. In particular, clonal expansion of TCR beta [13,27] and alpha [28] chains was demonstrated in AAA lesions of patients. In addition, there is a clonal expansion of gamma delta T cells [13] in AAA. Furthermore, TCRs were investigated in different types of TAA (patients with Marfan syndrome, familial thoracic aortic aneurysm and sporadic aneurysm) and the results indicate a similar clonal nature of the TCRs present in TAA [29]. He et al., found a preferential usage of the V-genes Vb22 and Vb25 in lesions from patients with thoracic aortic aneurysm [29]. Lu et al. reported multiple appearances (at least twice) of TRBV3 in 60% of AAA patients [27]. Atherosclerotic vascular disease, which is also a risk factor for AAA development [51], is likewise associated with T cell expansion and clonality. In particular, TCRs containing Vβ6 are expanded in atherosclerotic lesions of mice [52]. Moreover, a decreased diversity of the TCR β chain repertoire was shown in human atherosclerotic plaques due to expansion of a few T cell subclones [53].

Unlike human aneurysms, which are most likely to form over time due to persistent inflammation, the PPE model is characterized by rapid development of aneurysm. This distinction may explain the disparities in TCR and BCR clonality seen in our work. Since other studies found TCR clonality in AAA lesions of patients and the autoimmune disease hypothesis is still relevant, further investigation of TCR and BCR clonality is important.

The main limitation of our data set is the small number of lymphocytes leading to potential undersampling. The limited number of cells resulted from the naturally scarce source (i.e. minimal aneurysm size in mice), from additional sorting procedures, and not fully efficient sequencing. Indeed, we had to exclude many TCR and BCR sequences from the data due to inefficient sequencing. The undersampling leads to the issue that the copies of TCRs and BCRs in our data do not represent the real absolute number of copies present in AAA, and even the ratio of the various clonotypes to each other changes [50]. Therefore, biological interpretation of the data is limited. To overcome the problem of undersampling we suggest to focus on B or T cells and sort at least 5000 to 10000 cells. Our flow cytometer measurements revealed that on average 120 to 430 to B cells and 400 to 1600 T cells per mg AAA tissue can be isolated depending on the stage of AAA development. However, it is not possible to isolate this number of lymphocytes from control conditions (e.g., native or-sham-operated mice; on average 120-300 B cells, 400-500 T cells/mg aortic tissue). For the comparison of datasets from different researcher groups it is important to have a standardized workflow for sample preparation and the preprocessing of the data. Accordingly, we present an example and detailed workflow for the pre-processing steps in this paper.

## Conclusion

In summary, we found no evidence for clonal enrichment of T or B cells in experimental PPE-induced AAA. Our findings imply that for more precise characterization of TCR and BCR, receptor sequencing should take precedence over total RNA sequencing of individual cells. We recommend sorting B and T cells before sequencing to reduce contamination and the possibility of undersampling. This increases the likelihood of detecting rare TCR and BCR clones. Our advice would be to prioritize sequencing of the TCR and BCR libraries while also using Cellular Indexing of Transcriptomes and Epitopes by Sequencing (CITE-Seq) antibodies against B cells (CD19), T cells (CD3), and potentially subpopulations to better define the cells.

## Acknowledgements

We would like to acknowledge the assistance from Katarina Raba at the Core Flow Cytometry Facility at the Institute for Transplantation Diagnostics and Cell Therapeutics Düsseldorf. Computational infrastructure and support were provided by the Centre for Information and Media Technology at Heinrich Heine University Düsseldorf. We thank Tobias Lautwein for performing the single-cell sequencing and analyzing scRNAseq primary data.

Figures were created with Biorender.

## Funding

This study was supported by the following grants: Deutsche Forschungsgemeinschaft (DFG, German Research Foundation) - Grant No. 397484323 – CRC/TRR259; project A04 to H.W., project A05 to N.G.; We acknowledge the support of the Susanne-Bunnenberg-Stiftung at the Düsseldorf Heart Center.

## Conflict of interest

The authors declare no conflict of interest.

**Supplement Figure 1:**
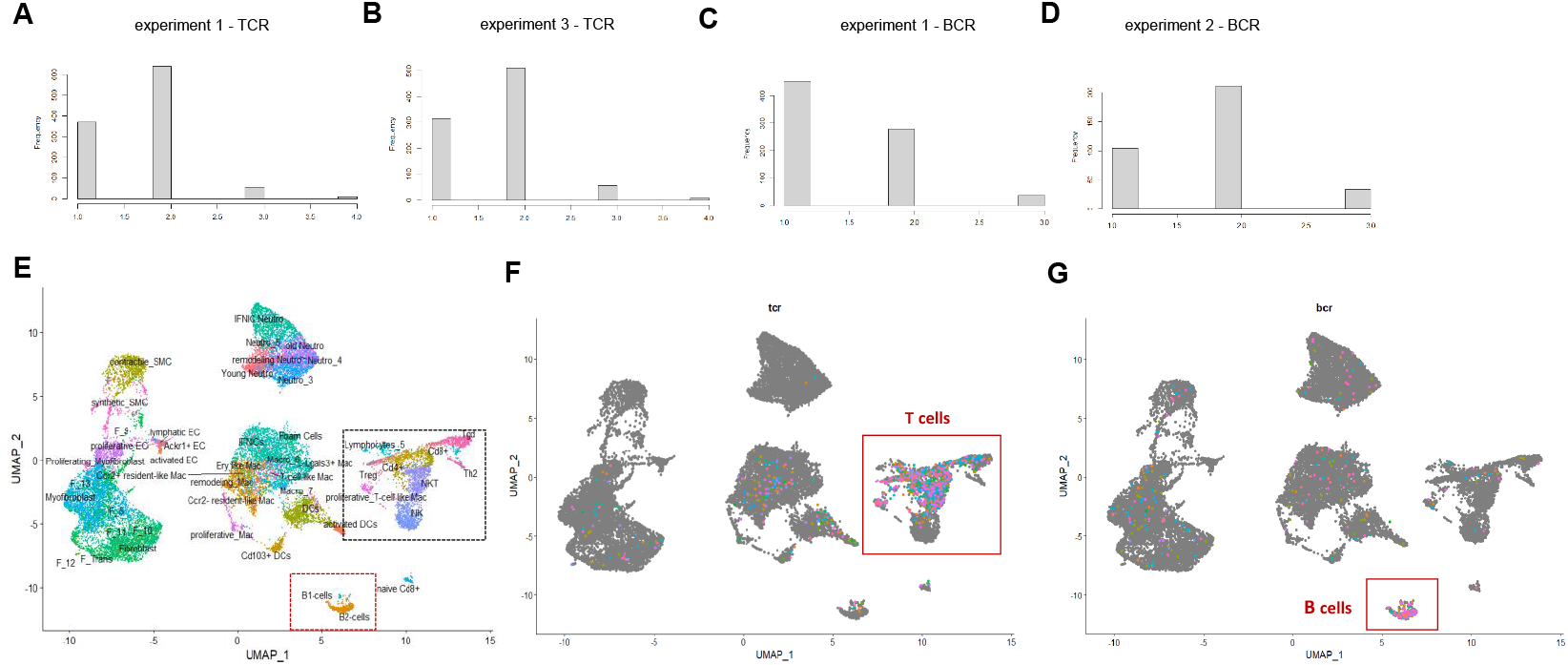
Preprocessing and quality control of scRNA TCR and BCR sequencing data. **A:** Number of chains per TCR in experiment 1. Only TCRs with two chains were used for analysis. **B:** Number of chains per TCR in experiment 2. Only TCRs with two chains were used for analysis. **C:** Number of chains per BCR in experiment 1. Only BCRs with two chains were used for analysis. **D:** Number of chains per BCR in experiment 2. Only BCRs with two chains were used for analysis. **E:** UMAP plot displaying all cell clusters in our scRNA sequencing data. T cell cluster are indicated with a black box. B cell cluster are indicated with a red box. **F:** Assignment of TCRs to the corresponding cells in scRNA sequencing data reveals that 79.13% of TCRs is expressed on T cells. Each TCR is indicated by a colored dot. **G:** Assignment of BCRs to the corresponding cells in scRNA sequencing data reveals that 46.49 % of BCRs is expressed on B cells. Each BCR is indicated by a colored dot.

**Supplement Figure 2:**
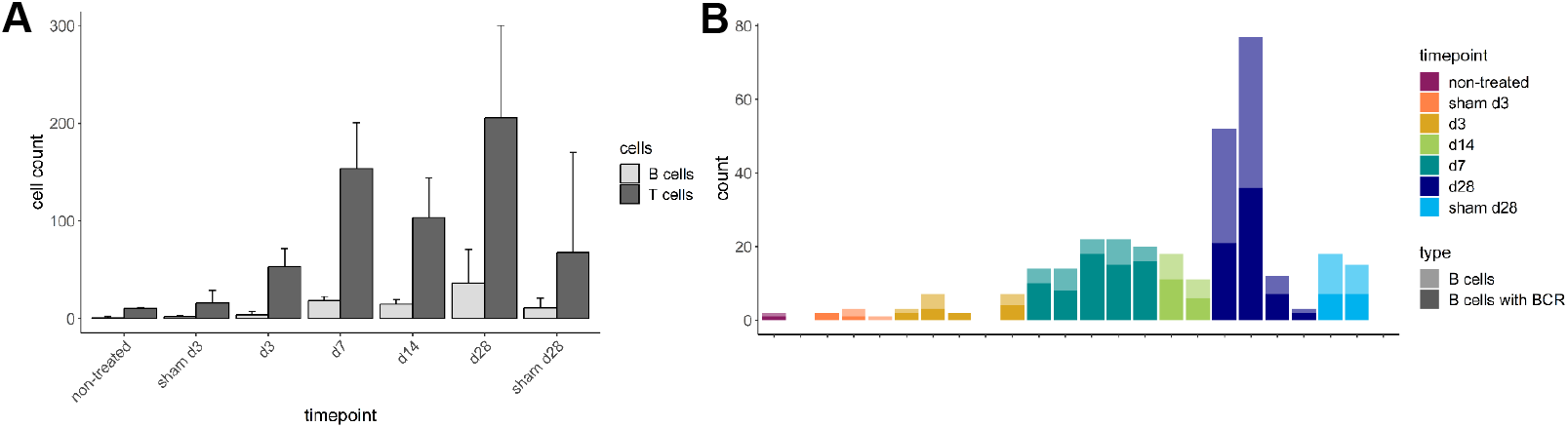
B cell numbers in AAA tissue increases over time. **A:** Number of B and T cells in scRNA sequencing data from non-treated, sham operated (d3 and d28) and PPE-perfused (d3, d7, d14 and d28) aortic tissue. Displayed is the mean ± SD. The amount of B and T cells increases with AAA progression, with the highest amount of cells on day 28 after PPE-induced AAA formation. **B:** B cell amounts (light bar color) and B cell amount exhibiting an intact BCR (dark bar color) in aortic tissue at different AAA disease stages received from scRNA Seq data. An intact BCR could be sequenced in 55.08% of the B cells.

**Supplement Figure 3:**
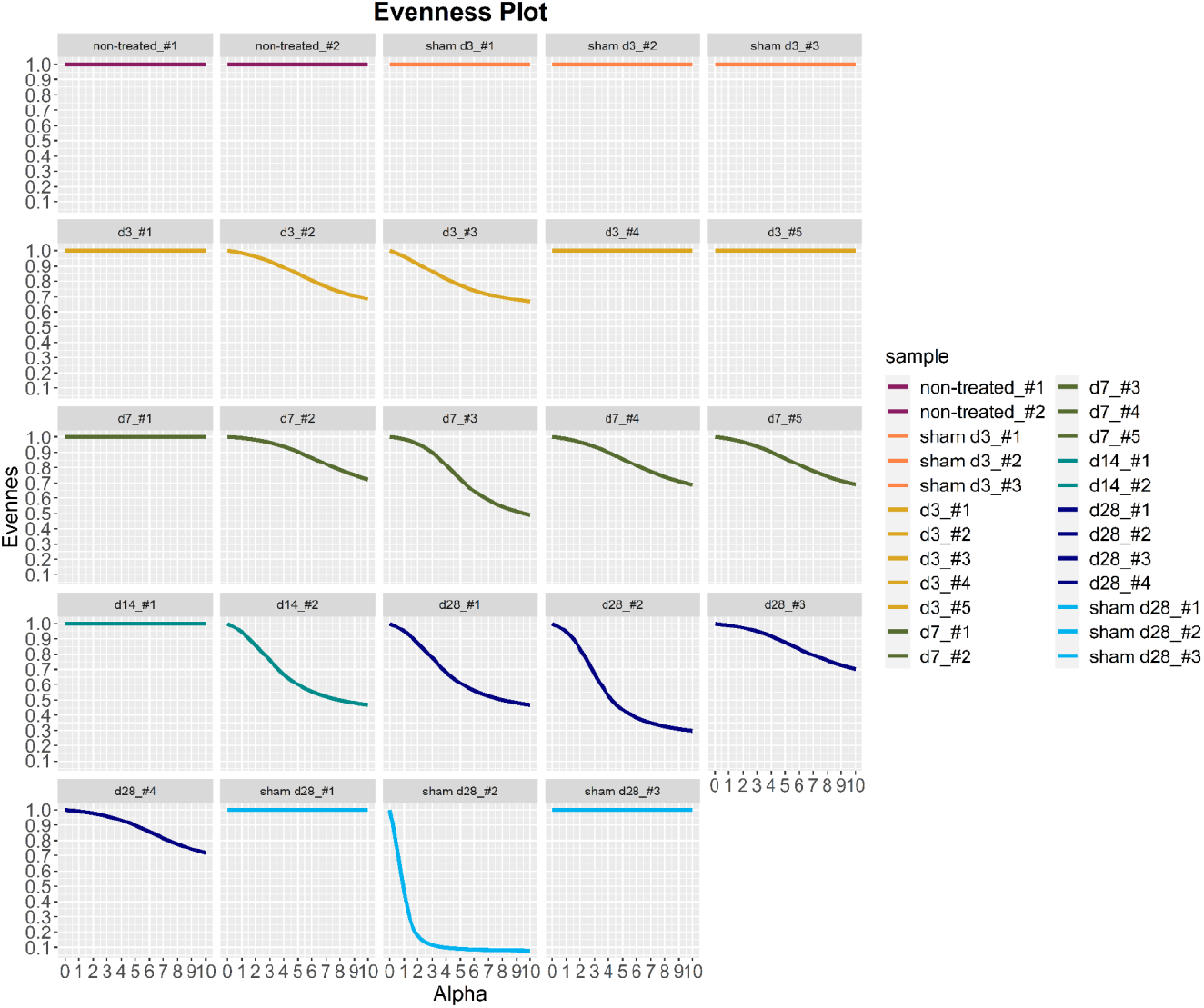
TCR receptor clonality was detected in 12 out of 24 samples. Evenness profiles of TCR repertoire showing the extent of clonal expansion for every sample. One sham operated sample 28 days after perfusion exhibit the highest clonality, whereas 12 samples show no receptor clonality. Eleven AAA samples show some clonal expansion.

**Supplement Figure 4:**
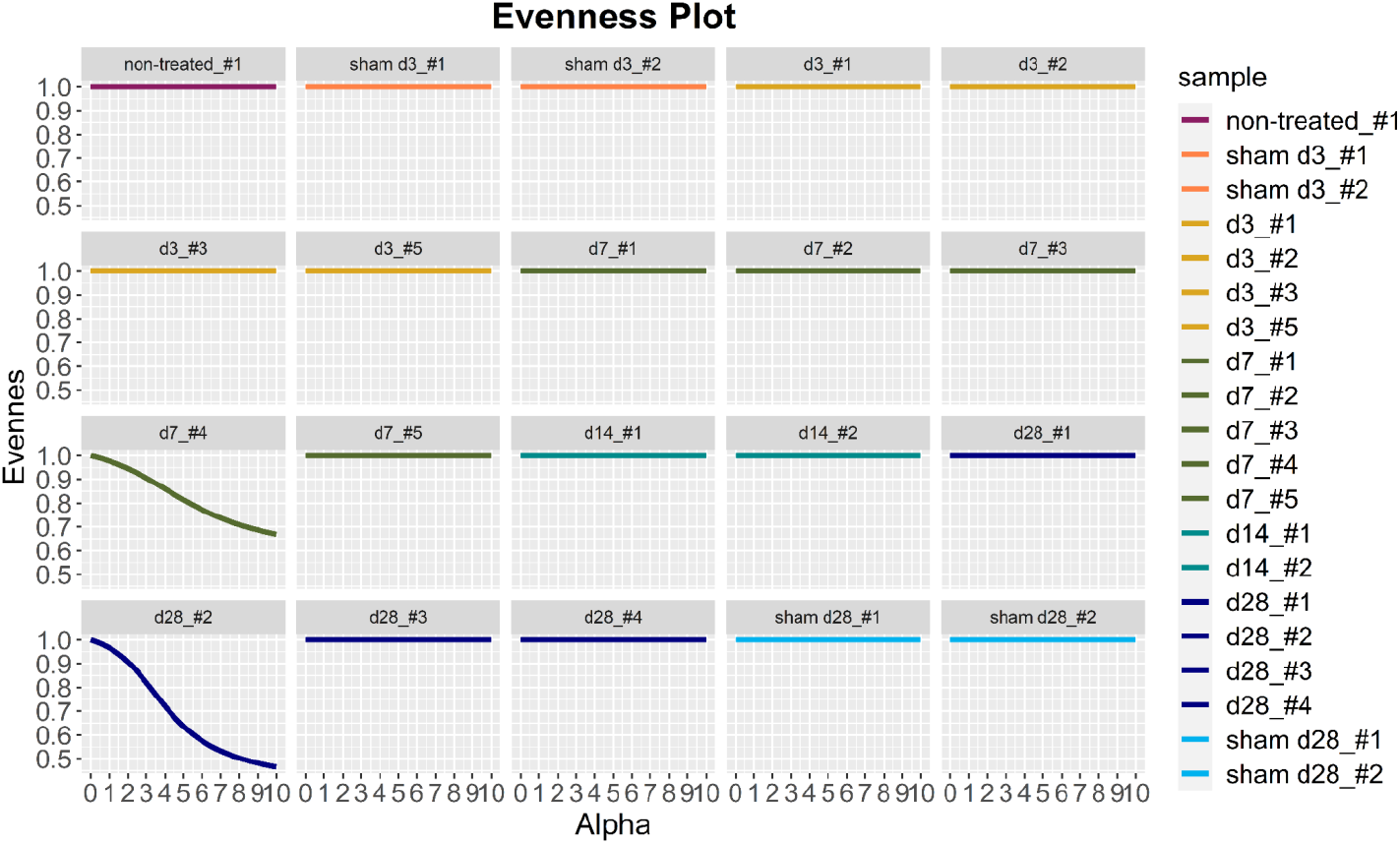
BCR receptor clonality was detected in 2 out of 20 samples. Evenness profiles of BCR repertoire showing the extent of clonal expansion for every sample. One AAA sample 7 days and one AAA sample 28 days after PPE-perfusion shows higher receptor clonality, whereas all other samples exhibit no clonal expansion.

## Notes

### Competing Interest Statement

The authors have declared no competing interest.

